# Docosahexaenoic Acid has stem cell-specific effects in the SVZ and restores olfactory neurogenesis and function in the aging brain

**DOI:** 10.1101/2020.02.10.942870

**Authors:** JE Le Belle, J Sperry, K Ludwig, NG Harris, MA Caldwell, HI Kornblum

## Abstract

Fatty acids are well known as important constituents for the synthesis of membrane lipids and as sources of cellular energy in the CNS. However, fatty acids can also act as vital second messenger molecules in the nervous system and regulate the activity of many proteins affecting cell growth and survival. Here, we show that an essential dietary fatty acid, Decosahexaenoic acid, (DHA), can enhance stem cell function in vitro and in vivo. We found that this effect is not due to an increase in the overall proliferation rate of all neural progenitors, but is due to an increase in the number of multipotent stem cells that leads to greater levels of subventricular zone (SVZ) neurogenesis with restoration of olfactory function in aged mice. These effects were likely mediated through increased EGF-receptor sensitivity, a conversion of EGRFR+ progenitors back into an EGRFR+/GFAP+ stem cell state, and the activation of the PI3K/AKT signaling pathway, which is a critical pathway in many NSC cell functions including cell growth and survival. Together these data demonstrate that neural stem cells in the aged and quiescent neurogenic niche of the mouse SVZ retain their ability to self-renew and contribute to neurogenesis when apparently rejuvenated by DHA and PI3K/AKT pathway activation. DHA stimulation of this signaling enhances the number of multipotent stem cells and neurogenesis in young and aged rodent and human stem cells and hence may have implications for the manipulation of neural stem cells for brain repair.

**Significance Statement:** We have identified potentially important effects of DHA on the stem cell population which may be unique to the SVZ stem cell niche. Our studies demonstrate that DHA can promote the production of neural stem cells, possibly via a non-proliferative mechanism stimulated by EGF receptor activation, and prolongs their viability. Aging animals undergo an apparent loss in SVZ stem cells and an associated decline in olfactory bulb function. We find that dietary DHA supplementation at least partially restores stem cell numbers, olfactory bulb neurogenesis and olfactory discrimination and memory in aged mice, demonstrating a capacity for rejuvenation is retained despite age-related changes to the niche, which has significant implications for ameliorating cognitive decline in aging and for endogenous brain repair.

## Introduction

The causes for aging-related cognitive decline are manifold. It has long been hypothesized that a large decrease in adult neurogenesis, which is known to occur with aging, is one factor that significantly contributes to this functional decline. In the adult mammalian brain, neurogenesis only occurs in specialized niche microenvironments like the forebrain subventricular zone (SVZ) and the subgranular zone (SGZ) of the dentate gyrus in the hippocampus (Bjornsson et al., 2015). It has been observed that adult SGZ neurogenesis is required for the plasticity of learning and memory (Drapeau et al., 2003). Similarly, SVZ-olfactory bulb neurogenesis is required for adult olfactory functions (Liu, et al., 2013).

Aging is associated with a decline in neural stem and progenitor cell proliferation and in the production of new neurons within both the SGZ and the SVZ (Ahlenius et al., 2009; Obernier et al., 2018; Shook et al., 2012; Silva-Vargas et al., 2016; Luo et al., 2006; Enwere et al., 2004; Maslov et al., 2004; Kalamakis, et al., 2019; Daynac et al., 2016; Bouab et al., 2011). In the SVZ, a focus of the current study, the functional consequences of this age-related decline are deficits in olfactory discrimination and olfactory memory (Gheusi, et al., 2000; Liu et al., 2013; Utsugi et al., 2014; Enwere et al., 2004; Rochefort, et al., 2002). Human studies of aging have shown that higher order cognitive processing is dependent upon odor recognition and memory, and that olfactory deficits parallel generalized age-related deficits in sensory functions and cognition that are seen with normal aging (Stevens & Cain, 1987; Chaker et al., 2015; Eichenbaum, et al., 2009). This depletion of the neurogenic capacity of the SVZ in aging may also have significant consequences for endogenous brain repair as SVZ-derived cells have been shown to migrate throughout the brain and produce new neurons in response to stroke and traumatic brain injury in rodent models (Lindvall and Kokaia, 2015). Though recent postmortem studies have called into question whether adult neurogenesis occurs to a significant degree from the SVZ in humans after birth (Kumar, et al., 2019), the potential restorative effects that DHA replenishment could have on this process have not been taken into account.

In contrast to other tissues, the nervous system is enriched in the polyunsaturated fatty acids: arachidonic acid (AA, 20:4 n-6) and docosahexaenoic acid (DHA, 22:6 n-3). Despite their abundance, neither AA nor DHA can be synthesized de novo by mammals due to a lack of the appropriate desaturase enzymes that introduce double bonds at the n-3 or n-6 carbon position required for their synthesis (see Figure 1A). Therefore, these essential fatty acids or their n-6 and n-3 precursors, linoleic and α-linolenic acid, must be ingested from dietary sources and transported into the brain. DHA is important for normal brain development (for review see McCann and Ames, 2005; Heird and Lapillonne, 2005) and its deficiency has been found to result in loss of visual and cognitive abilities in rodents and humans (reviewed in McCann & Ames, 2005 and Innis, 2003). Conversely, dietary supplementation of DHA during infancy may improve cognitive development in humans (Birch et al. 2000, Willatts et al., 1998) and may have preventative and ameliorative effects on cognitive function in aged, diseased (i.e., Alzheimer’s and Parkinson’s Diseases), or injured (i.e., stroke and traumatic brain injury) adult brain (reviewed in Johnson & Schaefer, 2006 and Uauy & Danquor, 2006; Arsenault et al., 2011; Calon, et al. 2004, Bousquet, et al., 2011, Belayev et al., 2009; Berressem et al., 2016, Bailes & Mills, 2010; Wu et al., 2011). DHA has been implicated in age-related brain function because declines in both brain and blood DHA levels have been correlated with the onset of dementia and cognitive decline (Weiser, et al., 2016; Mohajeri et al., 2015; Yurko-Mauro, et al., 2010). However, despite a considerable amount of compelling evidence regarding the interactions of DHA with brain function, the mechanisms and cell-specific effects which underlie its functional impact are still not fully understood.

**Figure 1.**
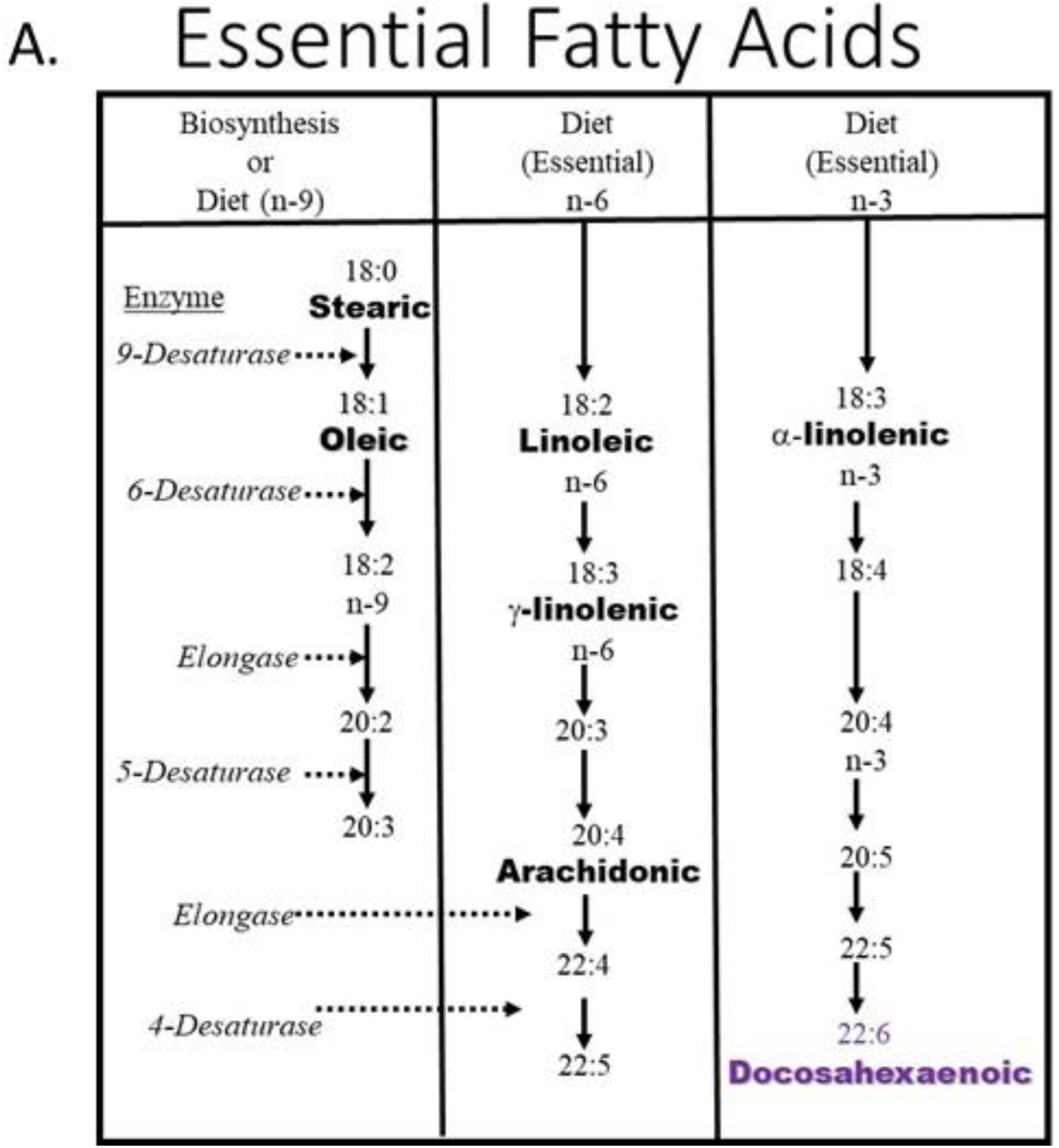
The essential fatty acid pathways diagram.

Here, we have examined the effects of DHA on neural stem and progenitor cells from the cortex of the developing brain and SVZ of the aged adult brain. We found DHA has the ability to significantly increase the number of neural stems and the neurogenic potential of the aged SVZ both in vitro and in vivo through a series of inter-related effects including increased sensitivity to EGF receptor activation, the trans-differentiation of a progenitor population, and elevated PI3K/AKT activity, resulting in a restoration of olfactory function.

## Materials and Methods

### Human and rodent neural cell culture

The cortical tissue was dissected from embryonic day 14 (E14) rat and mouse brain or from the subventricular zone (SVZ) of adult mice. Human embryonic cortical tissue (8 to 10 weeks post conception) was collected according to the specifications of the Polkinghorne Committee for the collection of human tissues and to the guidelines set out by the United Kingdom Department of Health. The brain tissue was acutely dissected and then treated with either trypsin (0.1% for 20 min; human and rat tissue) or Accumax (1X for 10 min; mouse tissue). Trypsin was inactivated using soybean trypsin inhibitor (Sigma) and washed in DMEM containing 0.05% DNase I (Sigma). After washing the cells were dissociated into a single cell suspension by gentle trituration with a fire-polished glass pipette. Cells were seeded at a density of 100,000 cells per ml in a defined, serum-free medium (DMEM:F12 at 2:1) supplemented with N2 (1% v/v; Invitrogen), 1mM L-glutamine, epidermal growth factor (EGF, 20ng/ml), and fibroblast growth factor (FGF-2, 20ng/ml) with heparin (5µg/ml). The serum-free supplement, B27, was not used in these experiments because it contains an undisclosed amount of the DHA precursor, linolenic acid, as well as L-carnitine, which would impact the examination of the effects of our DHA-supplemented conditions. Instead a similar serum-free supplement, N2, which does not contain essential fatty acid precursors or carnitine was used. Half of the growth medium was replenished every 4 days. The cells formed cell aggregates by 2 to 5 days of growth and were cultured as free-floating neurospheres according to the methods of Svendsen, et al., 1998. Briefly, the cells were passaged every 7-14 days by either sectioning the spheres into 150 µm sections (human cultures) or dissociating the spheres into a single cell suspension (human, mouse and rat cultures) and re-seeding into fresh growth medium. The following fatty acid supplementations were made to the standard culture media (all fatty acid-supplemented cultures also contained L-carnitine supplemented at 30uM and fatty acid-free bovine serum albumin at 1uM; all from Sigma): Docosahexaenoic acid (10 µM). The PI 3-kinase (PI3K) inhibitor LY294002 (Calbiochem) was used at 10µM. L-carnitine is an important fatty acid co-factor to transport long-chain fatty acids into the mitochondria for oxidative metabolism and modification which is primarily obtained via diet (Athanassakis, et al. 2002). Since it is not thought to be synthesized by cells from the developing brain (Arenas, et al., 1998), L-carnitine was supplemented with the fatty acids to facilitate their normal utilization in culture. Cells were grown from the time of dissection from the brain and dissociation into cell culture for three serial passages in either control or fatty acid-supplemented media before analysis in our experiments. The concentration range at which we found the fatty acids to have positive effects on cell survival, self-renewal, and neurogenesis is quite small (10-30 µM) and they are toxic to the cells at higher concentrations (data not shown). This is in agreement with the observations by others that have examined the role of DHA in neurogenesis and found a similarly effective window (1-10uM) in vitro before DHA has toxic effects (Insua et al., 2003; Kawakita et al., 2006).

### High density cell culture expansion

Cells grown in different media conditions were passaged by dissociation into a single cell suspension with trypsin and DNaseI as described above. Cells were re-suspended in 1 ml of culture media and a 10µl aliquot of live cells was counted on a haemocytometer for trypan blue dye exclusion. Total live cell content for each culture was calculated every 7 days for 8 weeks. After counting cells were re-seeded at high density (50,000 cells/ml) in one T25 flask per condition. At passage 3 sister cultures were grown separately for a further week before being plated down as whole-spheres for analysis of differentiation potential.

### Functional Serial Clonal Density Neurosphere-forming Assay

Neurospheres grown at high density (50,000 cells per ml) were dissociated into a single cell suspension as described above and re-seeded into 96-well plates at 20 cells per well in 200ul of media (100 cells per ml or clonal density). This density has been experimentally determined to produce single-color clonal spheres equivalent to single cell per well experiments by mixing red, green, and non-fluorescent cells at limiting dilutions (data not shown). Clonally-derived neurospheres were counted and their diameters measured using a brightfield illumination and image analysis software (MCID, Imaging Research, St. Catherines, ON, Canada). A minimum diameter cutoff of 35 µm was used in defining a neurosphere. Each plate contained culture media with growth factors and appropriately supplemented as previously described with one plate per condition. Clonally-derived neurospheres were counted and their diameters measured using a brightfield illumination and image analysis software (MCID, Imaging Research, St. Catherines, ON, Canada). A minimum diameter cutoff of 40 µm was used in defining a neurosphere. Spheres were then transferred onto poly-l-lysine coated chamberslides and differentiated for 5 days. Spheres were fixed with 4% paraformaldehyde for 20 minutes and immunostained for Tuj1, GFAP, and O4 as described under immunocytochemistry below. The number of multipotent neurospheres was then counted. Sister cultures at clonal density were maintained at the same time to passage and plate again in secondary and tertiary clonal cultures as described above. Repeated clonal passaging performed serially is used to establish true self-renewal capacity.

### Single-cell-per-well cell sorting

The isolation of stem cell-enriching antigen positive and negative cells for clonal analysis (Capela and Temple, 2002) was performed using Fluorescent Activated Cell Sorting (FACS). Sorted cells were either placed one cell per well directly in 96-well plates or collected in tubes depending on the experiment. Cells were labelled with antibodies to the stem-cell enrichment markers SSEA1 (Lex) (CamFolio, 1:200), EGFR (Cell Signaling Technology, 1:100), ID1 (Biocheck, 1:100), GFAP (Dako, 1:200) and the FAB7 dye (CDr3, a kind gift from Professor Chang Young-Tae; Yun, et al. 2012) for 30 minutes at room temperature with goat-anti-mouse Alexa 488, 548, 647, or PercP secondary antibodies (Molecular Probes) at a 1:2000 dilution for 30 minutes at room temperature. Background signal was determined by incubation of the same cells without primary antibody (negative control). FACS was performed with a FACSDiVa cell sorter (BD Biosciences) using a purification-mode algorithm. Sort gates were set by side and forward scatter to eliminate dead and aggregated cells and by secondary staining to define positive cells. Purity of the sorted cells was confirmed by flow cytometric re-analysis of positive and negative cell samples.

### BrdU labeling index

To label dividing cells with 5′-bromodeoxyuridine (BrdU), rat cortical cultures in all media conditions were incubated with 1μM BrdU for 4 h prior to dissociation and plating. Cells were dissociated with Trypsin as described above and plated on poly-l-lysine coated chamberslides for 1 hour. Cells were then fixed with 4% paraformaldehyde for 30 min and immunostained for BrdU label incorporation and counterstained with Hoescht as described below in the immunocytochemistry section. The percentage of total cells (Hoescht) immunopositive for BrdU was calculated for the 4-hour pulse-label.

### [3H]Thymidine incorporation

Human cortical cultures in all media conditions were incubated with [3H]thymidine (1 μCi/ml) for 12 h at 37°C. At the end of the incubation, cells were washed x3 with DMEM to remove the unlabelled 3[H]thymidine. This was followed by a further 3 washed with 10% trichloroacetic acid (TCA) and lysis with 0.1 M NaOH/0.1% sodium deoxycholate for 1 hour at 37oC. Lysates were transferred to vials containing scintillant and vigorously vortexed. Radioactivity was measured using a liquid scintillation spectrometer.

### Carbofluorescein-Succinimidyl-Ester (CFSE) dye wash-out

Murine cortical cultures in control or supplemented media were cultured for 4 passages before being dissociated into a single cell suspension and stained with 5 µM CFSE (Molecular Probes) in PBS at 37°C for 15 minutes in the dark. Cells were washed 3X in 5ml of HBSS-CMF. Immediately after staining, one half of the cells were fixed in 1% paraformaldehyde and analyzed in a Becton Dickenson FacsCaliber flow cytometer. The remaining cells were cultured under neurosphere culture condition (see above). After 4 days, neurospheres were dissociated and fluorescent intensities were measured by flow cytometry as above. As a control, 10µM Aphidicolin (Sigma), an S-phase cell cycle blocker, was added to one of the samples. (n=2).

### Cell Titer Blue Assay

Cell viability was measured using a fluorometric assay kit (Promega Celltiter Blue cell viability assay). Murine cortical cultures in control or FA-supplemented media for 4 passages were dissociated into a single cell suspension and seeded at a density of 5,000 cells per ml in 100 µl media in 96-well plates and the viability assay performed 24 hours later by adding 20 uL CellTiter Blue reagent followed by a six-hour incubation at 37°C. Data was collected on an Analyst HT (Molecular Devices, Sunnyvale CA).

### Caspase 3/7 activity assay

Caspase activity was measured using a chemiluminescent assay kit (Promega Caspase-Glo^®^ 3/7) in 96-well format using an Analyst HT plate reader (Molecular Devices, Sunnyvale CA). Murine cortical cultures in control or DHA-supplemented media for 4 passages were dissociated into a single cell suspension and seeded at a density of 5,000 cells per ml in 100µl of media and the caspase assay performed 24 hours later according to kit protocol.

### Cell Differentiation potential

Intact neurospheres were plated down on poly-D-lysine- (100 µg/ml) and laminin-coated (10 µg/ml) chamberslides. Cell mitogens (EGF & FGF) were withdrawn from the different cell media compositions and the plated cells were allowed to differentiate for 5-7 days before being fixed for immunostaining.

### Immunocytochemistry

Cells were fixed in 4% paraformaldehyde in PBS containing 4% sucrose for 20 min. Immunostaining was carried out using standard protocols. For BrdU staining, cells were treated with 2 M HCl for 20 min at 37°C followed by two 5 min washes in 0.1 M sodium borate buffer. 3% normal goat serum was used to block all cells before addition of primary antibodies. Primary antibodies and dilutions were as follows: β-tubulin type III monoclonal (TuJ1, 1:500; Sigma, St. Louis, MO), GFAP polyclonal (1:1,000; DAKO, Glostrup, Denmark), GFAP monoclonal (1:500; Chemicon, Harrow, UK), TH monoclonal (1:500; Chemicon), BrdU monoclonal (1:300; Roche, Sussex, UK), GABA polyclonal (1:250; Sigma), O4 (1:50 hybridoma supernatant; gift from De Vellis lab). Cells were then reacted with appropriate secondary antibodies (Alexa 488 or 568, Molecular Probes); 1:500) for 1 h at room temperature. Hoescht nuclear stain (1:5000) was included in the final antibody applications. In order to determine the number of neurons and astrocytes, and oligodendrocytes photographs of immunostained cells were taken using a high-resolution digital camera (Nikon). Five fields per chamber were counted with eight chambers per condition in each experiment.

### Western Blotting

All primary antibodies (total Akt and phospho-specific Akt) and positive and negative controls were purchased from Cell Signaling Technologies. Neurospheres from each condition were lysed in buffer containing 0.1% triton X-100 in 50 mM Tris-HCl and 150 mM NaCl and Protease Inhibitor Cocktail (Sigma). Samples were then sonicated and subsequently spun down at 13,000 RCF for 10 minutes at 4°C. Supernatants were separated, and protein concentrations were determined by Bradford assay using BSA as a standard. 20ug of protein from each sample was separated on a 4-20% gradient Tris-HCl denaturing gel by SDS-PAGE, transferred to PVDF membranes for 1 hour and blocked with 5% non-fat milk. The membrane was incubated with primary antibodies overnight at 4°C followed by a series of 3 washes in PBS. Membranes were then incubated with a HRP-conjugated Goat anti-Rabbit secondary antibody for 1 hour at room temperature followed by 3 washes in PBS. HRP detection was performed using the ECL Plus (Thermo Fisher) and membranes were imaged using a Sapphire Biomolecular Imager (Azure).

### *In vivo* Dietary DHA Supplementation

Algae-generated DHA oil (DHASCO) was a generous gift from Martek Biosciences (USA) which was used by Harlan Teklad (USA) to produce custom rodent feed either enriched in DHA (28 mg/kg; TD.08416) or deficient in DHA (control diet: Global 16% protein; TD.00217). Young adult mice on the control diet were used to establish the baseline of normal adult SVZ proliferation and neurogenesis. These mice were fed the control diet for 6 weeks. Aged mice (1.4 yrs old) were divided into two groups and fed for 6 weeks on either the DHA-enriched or control diet. After 1 week the mice were injected with BrdU (50 mg/kg) once per day for 7 days to label all dividing cells. The mice continued to be fed their respective diet for a further 4 weeks during which time labelled cells were able to migrate and differentiate into new neurons in the brain. Mice were tested for olfactory function in two behavioral tests prior to being perfused for brain immunohistochemistry analysis or dissected for culture or flow cytometry analyses. Brains were prepared for immunostaining as described above. Antibodies for cells that were actively dividing at the time of perfusion (Ki67), for the BrdU label (Exalpha), and for differentiated neurons (NeuN, Abcam 1:200) were used to stain the tissue before sterological quantification of cell number in the SVZ or olfactory bulb was performed.

### Stereological cell counting

The estimated total number of labeled cells was determined by epifluorescence microscopy using unbiased stereology cell counts with the optical fractionator method, as implemented by StereoInvestigator software (MicroBrightfield, Williston, VT, USA) in 6, 20 µm sections within the SVZ region beginning rostrally at the genu of the corpus callosum and within the olfactory bulbs. Labeled cells were counted at 40x total magnification using a grid size of 35 × 35 µm with a sampling box of 15 × 15 µm and were reported as estimated cell populations.

### Olfactory Behavioral Tests

Examination of the relative ability of the mice to effectively use olfaction in two behavioral tests was performed in aged mice fed either the DHA-enriched or DHA-deficient diet and in young adult (8-10 weeks old) mice used as a baseline control for normal olfactory function. The first test examines the natural mouse behavioral preference for social novelty which requires the use of the sense of smell to distinguish between familiar and stranger mice (Bluthe et al., 1993; Lüscher Dias, et al., 2016). This test is commonly referred to as the three-chambered social approach task for mice where the mouse can freely move between 3 chambers containing either nothing (center chamber), a novel object (empty mesh or slatted cup) or a novel mouse in a mesh or slatted cup (Moy et al., 2004). In the case where this testing paradigm is used to test for olfactory based memory function, this phase of test is a training phase where the subject mouse under normal circumstances spends more time in the chamber containing the novel mouse than in the chamber containing the inanimate novel object or the empty chamber. All of our mice tested performed normally during this training phase. The second phase of this test takes place the following day when one chamber contains in a slatted cup the familiar mouse that it interacted with previously but the other, opposite chamber now contains a novel mouse in a slatted cup. Normal social recognition and social novelty seeking in this phase of the test is defined as the subject mouse spending more time in the chamber containing the novel mouse than in the chamber with the familiar mouse. Olfaction plays a key role in social recognition in rodents, since either chemically induced anosmia or removal of the vomeronasal organ blocks individual recognition (Matochik, 1988, Bluthe et al., 1993). The second test of olfactory function is the buried food detection test (Yang and Crawley, 2009), which relies on the animal’s natural tendency to use olfactory cues for foraging. The latency for a mouse to uncover a small piece of palatable food (Honey Teddy Grahams, Nabisco), hidden beneath a 2-inch layer of clean cage bedding, within a limited amount of time is measured and compared between groups. Mice are food-restricted overnight before this test to motivate foraging. Thus a failure for food-restricted mice to use odor cues to locate the food within a 15 minute period are likely to have deficits in olfactory abilities. The majority of mice with normal olfaction can find the hidden food piece within a few minutes.

### EGF trans-differentiation test

Adult mouse brains were sectioned coronally in 1mm sections and the SVZ region was micro-dissected from slices. Dissected tissue was enzymatically digested using Accumax (Sigma-Aldrich, USA) and mechanically triturated using a fire-polished Pasteur pipette into a single cell suspension which was passed through a 40um nylon mesh. The acutely dissociated SVZ cells from 5 mice were pooled and then divided evenly into 4 groups (control, DHA only, EGF only, or DHA + EGF) and placed in culture media either with or without 10 uM DHA and with or without 20ng/ml EGF for a 4 hours incubation. Following these acute treatments the cells were all then immunostained with an cell surface-EGFR antibody conjugated with FITC secondary antibody (Cell Signaling Technology) in Hibernate media (BrainBits) at room temperature for 30 minutes. After the EGFR staining, the cells were fixed and permeabilized using the Fix & PERM kit (Invitrogen) and immunostained with polyclonal GFAP (Gibco) and anti-rabbit PerCP secondary (Invitrogen) according to manufacturer instructions. The stained cells were analyzed using a FACSDiVa cell sorter (BD Biosciences) flow cytometer and FlowJo analysis software (Treestar). Sort gates were set by side and forward scatter to eliminate dead and aggregated cells and with negative, unstained control cell samples.

### Statistical analysis

Statistical comparisons were made by one- and two-way ANOVA, where appropriate, with post hoc Bonferroni t-tests (Sigma Stat software). Data are expressed as mean + standard error of the mean (SEM).

## Results

### Dietary DHA rescues age-related declines in SVZ proliferation, NSC number, and olfactory neurogenesis in vivo

Given the significant clinical effects of dietary DHA supplementation that have been observed in humans on brain function in aging in humans, we wanted to determine if DHA can restore the age-related declines in SVZ and OB neurogenesis and function that are known to also occur mice. We placed aged mice (73 weeks) on a DHA-enriched diet or DHA-deficient control diet for 1 week before labelling the dividing cells in the SVZ with daily pulse of BrdU injections for 1 week, followed by a 4 week maturation period for the labelled cells (see experimental paradigm in figure 2A). We found that the DHA-enriched diet produced a significant increase in the number of actively proliferating Ki67+ cells within the SVZ at the time of euthanasia (week 6 of DHA diet), in the number of BrdU+ cells that migrated to the olfactory bulb during the 4 weeks after labelling, and in the number of BrdU+/NeuN+ double labelled, newly-generated neurons within the olfactory bulb (Figure 2B). Similarly, acute dissections and cell dissociation from the SVZ of aged mice consuming a DHA-enriched or control diet and young adult mice consuming a control diet were used in the ex vivo serial clonal density functional culture assay to examine the number of self-renewing neural stem cells that were present in the SVZ from each group. We found that while there is a large loss of neural stem cells in the SVZ due to age, the 6 weeks of treatment with the DHA-enriched diet was able to significantly increased their number (figure 2C). These findings suggest that the DHA diet can increase the number of SVZ neural stem cells, the number of proliferating SVZ cells, and neurogenesis in the olfactory bulb in aged mice in vivo, though there is not a complete rescue back to the levels seen in the young adult SVZ.

**Figure 2.**
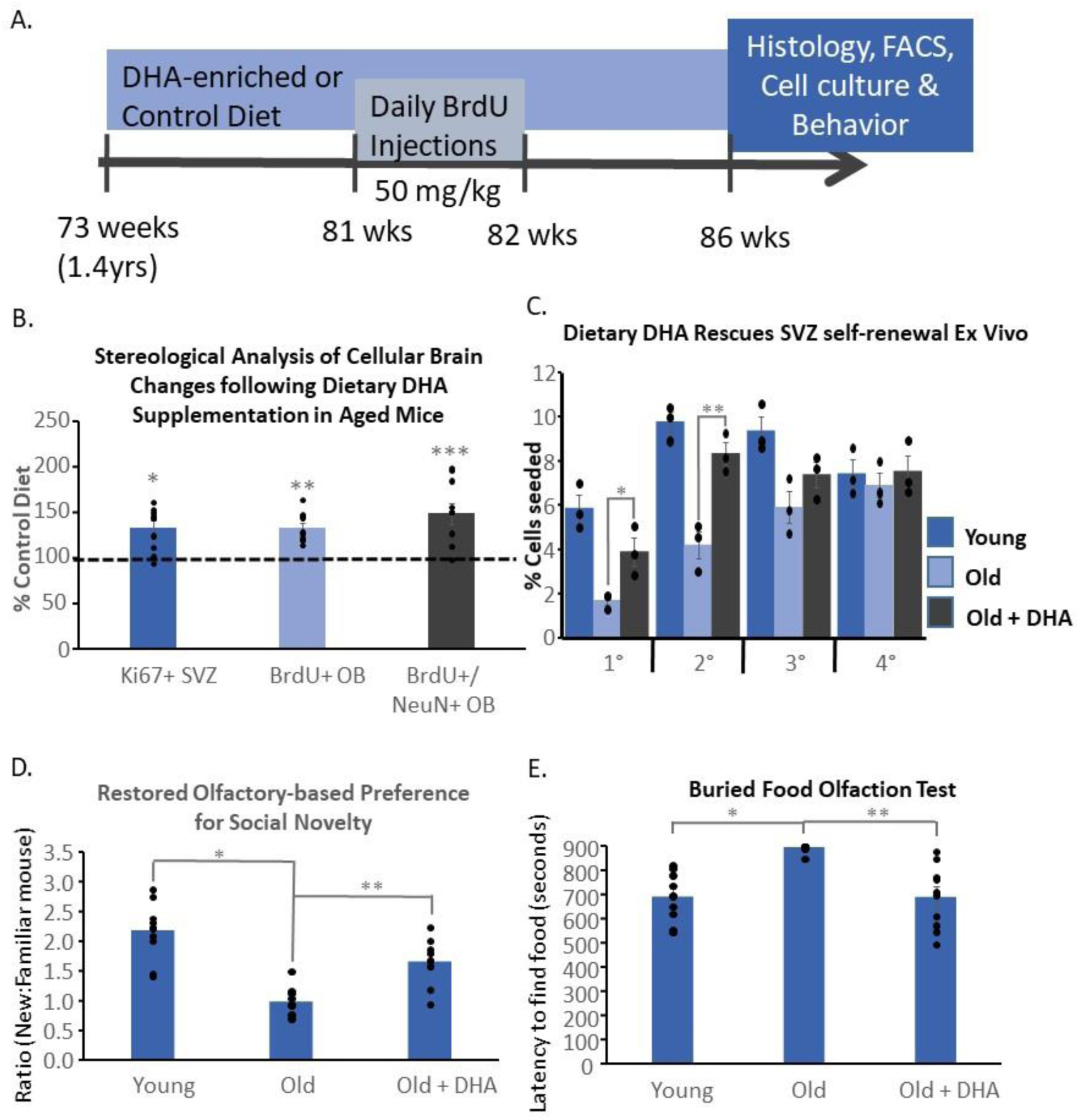
Dietary DHA enrichment enhances *in vivo* SVZ function in the aged mouse brain. (**A**) Experimental timeline demonstrating mice aged 18 months (old mice) were placed on a DHA-enriched or -deficient diet for 3 months, one week of BrdU injections were given to label dividing cells, and 1 month later the mice were sacrificed and their brains were processed for histological examination of the numbers of new neurons produced in the olfactory bulb and proliferating cells in the subventricular zone (SVZ) of the lateral ventricles; (**B**) Stereological counting of cells in the SVZ and olfactory bub indicate that an *in vivo* DHA-enriched diet significantly (P<0.001 ANOVA Bonferroni post hoc *p=0.024,**p=0.019, ***p=0.0001; n=10/grp) increases proliferation in the SVZ (Ki67+) and the migration and maturation of newly born neurons found in the olfactory bulb (BrdU+/NeuN+) in old mice compared to old mice fed a control diet; (**C**) Mice treated *in vivo* under the same dietary regimen described above were used to generate clonal density, multipotent neurosphere cultures from acute SVZ dissections which demonstrated a significantly increased number of stem cells (P<0.001 ANOVA; Bonferroni post hoc *p=0.0305, **p=3 ×10^−3^, n=6/grp) present in the SVZ of old mice fed the DHA-enriched diet compared to old mice fed the control diet, though the number is not fully restored to the levels seen from the young (8-10 weeks old) brain. Four serial clonal density passages are represented (1°-4°); (**D**) Behavioral testing for olfactory function indicates that old mice have a deficit in forming olfactory-based social memory in the 3-chamber social interaction test that is significantly (P>0.001 ANOVA; Bonferroni post hoc *p=0.0004, *p=0.001 n=10/grp) rescued by the DHA-enriched diet; (**E**) Behavioral testing for olfactory sensitivity indicates that old mice have a deficit in their ability to identify buried food treats within the 15 minute testing period through olfactory detection which is also significantly (P>0.001 ANOVA; Bonferroni post hoc *p=4×10^−6^, **p=1×10^−4^ n=10/grp) rescued by the DHA-enriched diet. Data is presented as mean +/- SEM.

### The DHA enriched diet rescues olfactory function in aged mice

We next examined if the significant restoration in olfactory bulb (OB) neurogenesis that we observed in the aged mice consuming the DHA-enriched diet were enough to affect OB function. At the end of the 6 week treatment period we examined olfactory function in aged mice fed the DHA or control diets compared to the olfactory function in young adult mice using two different olfactory behavioral tasks. The first test is a social memory task based on the ability to identify a novel versus a familiar mouse on the basis of smell. Using a standard 3-chamber social test, the mice freely move and choose to spend time in the different chambers: one containing a stranger mouse, one empty, and one containing an inanimate object. All of the mice performed normally in this training period, showing a strong preference for spending time in the chamber with the mouse over the other chambers (data not shown). The next the day the olfactory-based social memory is tested by measuring the amount of time the subject mouse spends in the chamber containing the familiar mouse from the previous day or in another chamber containing a novel mouse that hasn’t been experienced before. The young adult mice performed normally in this task, showing a strong, normal preference for the novel mouse over the familiar mouse and the empty chamber (Figure 2D). The aged mice on the control diet showed a clear deficit in distinguishing between the familiar and novel mice in the test but the old mice that were fed the DHA-enriched diet were significantly improved, showing a normal preference for social novelty based on olfactory-based discrimination between the familiar and novel mice (Figure 2D). We then performed a second olfactory test of the ability of the mice to detect the presence of desirable food buried under standard cage bedding. The latency for each mouse to find the buried food treat was measured with the test ending after 15 minutes. The younger adult mice performed normally on this task, finding the buried food within 11 minutes on average, but the aged mice were significantly impaired with most not locating the food at all. However, the aged mice fed the DHA-enriched diet located the food rewards as quickly as young adult mice, showing significant improvement in olfaction in this test as well (Figure 2E).

### In vitro experiments indicate DHA increases neural stem cell self-renewal but doesn’t significantly affect the overall proliferation rate of neural progenitors

Neurospheres are complex cultures, consisting of stem cells, progenitors, and a small number of differentiated cells. In order to determine what the cellular substrates for the effects of DHA on these cell populations, we performed a functional serial clonal neurosphere forming assay which measures stem cell self-renewal (serial clonal sphere number), overall proliferation (sphere diameter), and multipotency (differentiation potential). We tested these cellular functions on embryonic rat and mouse cortical cultures, adult mouse subventricular zone-derived cultures, and human fetal cortical cultures. The cells were seeded at a density such that all spheres would be derived from single cells (clonal density; Le Belle et al., 2011). Results show that the in vitro DHA-supplemented cultures had large increases in multipotent neurosphere-forming cells compared to controls but there was no significant effect on the overall proliferation rate (diameter of spheres) or in the number of clonal neurospheres that were multipotent (number containing neurons, astrocytes, and oligodendrocytes) (Figure 3A). This lack of effect on stem and progenitor proliferation rate was further confirmed using multiple different cell proliferation assays (BrdU labelling index, 3[H]thymidine uptake, and CFSE dye-washout; Figure 3B). Further examination of the differentiation potential beyond just multipotency of the DHA treated cultures demonstrated an enhanced neurogenic potential following growth factor withdrawal (Figure 3C). Although glial differentiation was not altered, the number of cells remaining undifferentiated in the DHA cultures was slightly decreased (figure 3C). These results suggest that our in vivo observations of increased numbers of proliferating cells in the SVZ and OB neurogenesis in animals given the DHA-enriched diet may not have been the simple pro-proliferative effect that we initially thought was underlying the increased SVZ neurogenesis. Indeed, our in vitro results suggest that it is the rescue of the number of neural stem cells present in the SVZ that restores the populations of proliferating progenitors and neuroblasts in vivo rather than simply enhancing the rate of stem and progenitor proliferation and neurogenesis.

**Figure 3.**
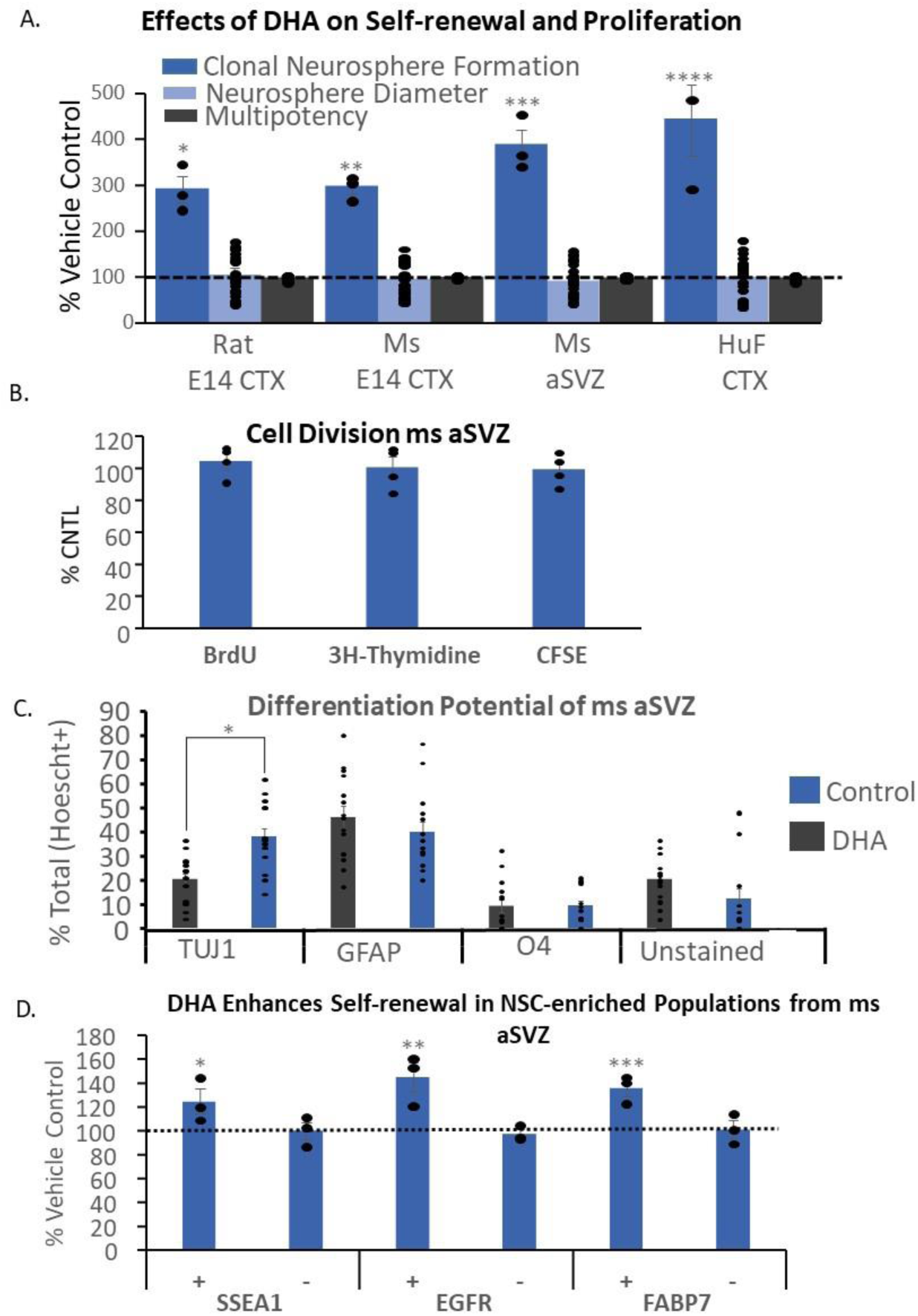
DHA supplementation in vitro increases neural stem cell numbers and neurogenesis but does not increase neural progenitor cell proliferation overall. (**A**) Embryonic cortex and adult SVZ neurosphere cultures generated from rats, mice, and humans have significantly more (P<0.0001 ANOVA, post hoc *p=4.67×10^−6^, **p= 1.2×10^−6^, ***p=7.9×10^−8^, ****p= 1.18×10^−6^ n=3/grp sphere number, n=10/grp diameter, n=15/grp multipotency) multipotent neural stem cells (clonal neurosphere numbers) than vehicle-treated cultures but there is no increase in overall proliferation (diameter) or multipotency; (**B**) Multiple measures of cellular proliferation (thymidine and BrdU uptake into dividing cells and CFSE dye wash-out) in adult mouse SVZ cultures indicate that there is no increase stimulated by DHA supplementation *in vitro* (P=0.925, n=4/grp) (**C**) Differentiation of intact clonally derived adult mouse SVZ neurospheres demonstrates that DHA supplementation of cultures significantly increases neuron differentiation (*p<0.001 Anova, Bonferroni post hoc *p=0.027;n=15/grp) without affecting glial differentiation; (**D**) DHA supplementation of adult mouse SVZ neurosphere cultures significantly (*p<0.0001 Anova, Bonferroni post hoc *p=3.72×10^−7^, **p=1.28×10^−9^, ***p=7.84×10^−11^; n=3 samples/grp; 3 mice were pooled per sample) increases multipotent clonal density neurosphere formation only in cultures that have been enriched in neural stem cells through FACS selection for SSEA1, EGRF, and FABP7, which are well-known stem cell enriching epitopes. DHA does not enhance stem cell numbers in the negative population of cells depleted of neural stem cells. All cultures were supplemented with DHA or vehicle (1% DMSO) from the time the cultures were established. Each culture spent a minimum of 4 weeks in these conditions. Data shown are mean +/- SEM.

### DHA effects on neural cultures are stem-cell specific

Because the in vivo effects on DHA appeared to affect the stem cell population specifically we next sought to further test if that is the case by examining the effects of DHA on cultures enriched for or depleted of neural stem cells through FACS sorting for known stem cell-enriching cell surface markers. Murine cells were live-sorted as a single cell per well in 96 well plates for Lewis-X, EGFR, or FABP-7. These antigens have been shown to enrich for NSCs, to be highly expressed in the cortex during development and to persist in the SVZ of the adult (Yun et al., 2012; Leong, et al., 2013; Capela and Temple, 2002; Pastrana, et al., 2009). Our results show that DHA supplementation enhances the number of multipotent clonal neurospheres (stem cells) to a greater extent within the stem cell-enriched cell cultures and had little to no effect on the cultures from the negative (stem cell-depleted) fraction of cells (Figure 3D), suggesting a stem-cell specific effect of DHA underlying the increases in stem cell numbers and neurogenesis in the SVZ of the aged mice in vivo.

### DHA increases NSC sensitivity to EGF stimulation

Given the importance of EGF signaling and EGFR expression in neural stem and progenitor cell populations and the possibility of altered membrane fluidity by DHA affecting receptor expression (Rapoport 2001; Turk, et al., 2012; Sinclair, 2019), we tested the effect of the DHA-enriched diet on the sensitivity of stem cell self-renewal to EGF stimulation. We found that cells freshly dissociated from the SVZ of the aged mice consuming the DHA-enriched diet in vivo produced more multipotent clonal neurospheres in response to stimulation by increasing EGF concentrations compared to the freshly isolated SVZ cells from aged mice consuming the control DHA-depleted diet, indicating an elevation in sensitivity to receptor stimulation as a result of the in vivo dietary DHA intake (Figure 4A). Therefore, we also aimed to determine if in vitro DHA treatment also alters the sensitivity of the SVZ cells to EGF activation. To do this we cultured EGFR+ FACS-selected cultures in a range of EGF concentrations and measured clonal neurosphere formation under those conditions. Unsurprisingly, cultures with no growth factor formed very few spheres (Figure 4B) and sphere formation increased with increasing EGF concentration but this effect was amplified in the cells that had been cultured with DHA (figure 4B). We then used FACS to quantify both the number of cells expressing the EGF receptor (EGFR) and the number expressing phospho-EGFR (activated EGFR) in cultures grown with or without DHA supplementation in the presence of normal culture levels of EGF. We observed a slight decrease in the expression of (cell surface) EGFR in the DHA cultures but an increase in EGF receptor activity (phospho-EGFR; Figure 4C), indicating an increased sensitivity to receptor stimulation by EGF.

**Figure 4.**
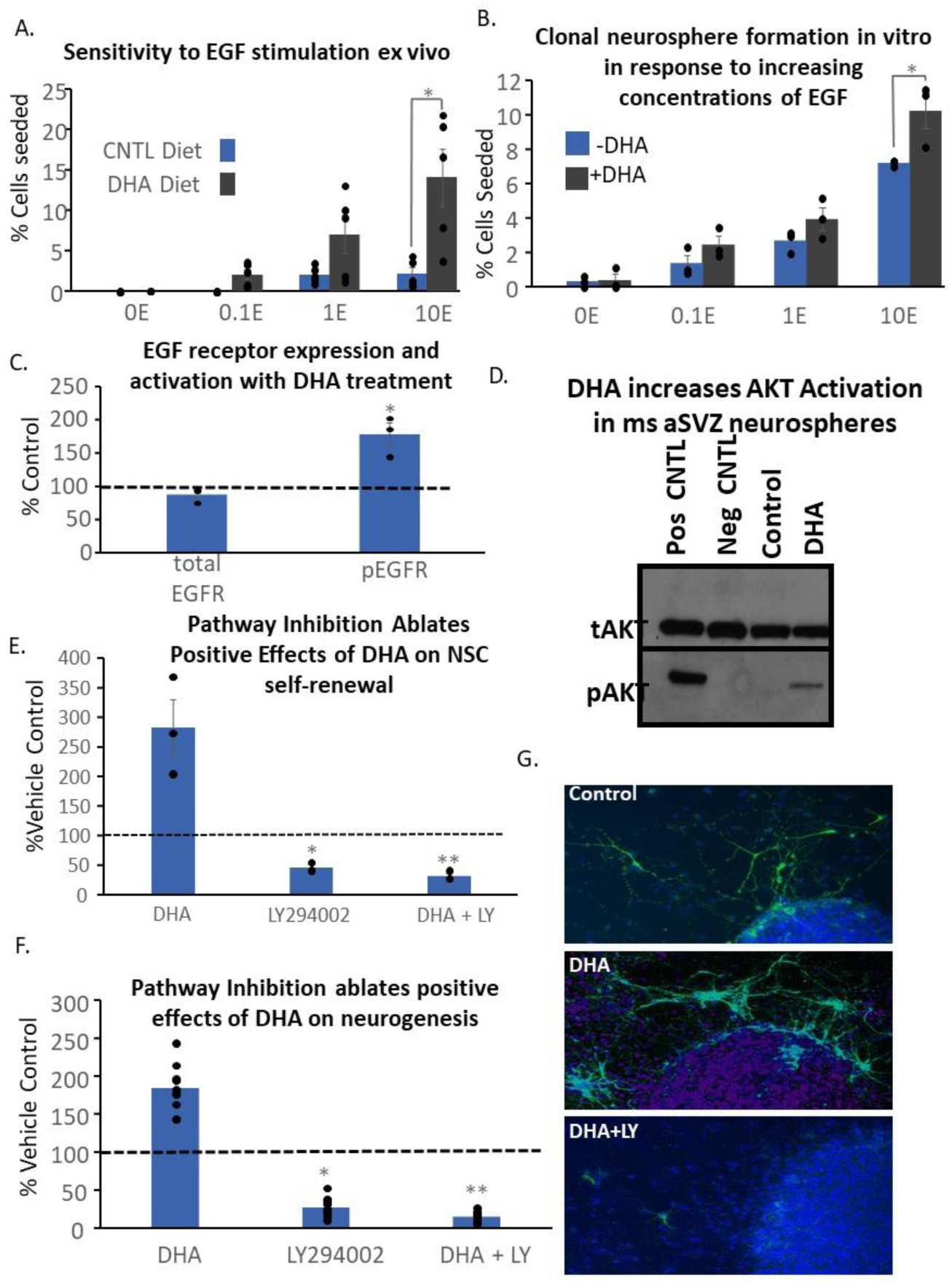
The effects of DHA on neural stem cells occurs largely through activation of the PI3K signaling pathway and increased sensitivity to EGF stimulation. (**A**) Old mice fed the DHA-enriched diet or control diet as described in Figure 1 were used to examine the sensitivity to EGF stimulation in acutely dissected SVZ cells that were FAC-sorted for EGFR-positive cells. No clonal neurospheres were formed without any EGF supplementation but the DHA-fed mice formed more clonal neurospheres with increasing EGF concentrations (P<0.001 Anova, post hoc Bonferroni *p=0.0106; n=5/grp); (**B**) Adult mouse SVZ cells also display increased clonal density neurosphere formation with increasing EGF concentration when supplemented with DHA *in vitro* compared to vehicle control cultures; (P<0.001 Anova, Bonferroni post hoc *p=0.038, n=3/grp (**C**) Adult mouse SVZ neurosphere cultures cultured with DHA over 5 passages have decreased but n.s. total EFGR expression but more (P=0.004 Anova, bonferroni post hoc *p=0.018, n=3/grp) phospho-EGFR expression after being starved of EGF for 12 hours and stimulated for 10 minutes before being fixed and permeabilzied for FACS acquisition analysis compared to the cells in control conditions (n=3/grp); (**D**) Representative western blot showing increase phospho-AKT expression in adult mouse SVZ neurosphere cultures supplemented with DHA compared to cultures in vehicle control conditions; (**E**) the PI3K inhibitor LY294002 significantly inhibits clonal density multipotent neurosphere formation in adult mouse SVZ cultures and similarly inhibits DHA supplemented cultures (P<0.001, Anova post hoc Bonferroni *p=0.0189, **p=1.06×10^−3^, ***p=4.47×10^−4^; n=4/grp) indicating the positive effects of DHA are not still produced by other signaling pathways; (**F**) LY294002 also significantly inhibits neuron differentiation from clonal density adult mouse neurospheres both with and without DHA supplementation (P<0.001 Anova, post hoc Bonferroni *p=4.94×10^−8^, **p=7.83×10^−10^; n=10/grp) (**G**) Representative immunocytochemistry demonstrating the effects of PI3K pathway inhibition on neuron differentiation from clonal adult mouse neurosphere cultures; Data shown are mean +/- SEM.

### DHA supplementation enhances PI3K/AKT pathway activation

Both EGFR activation and neural stem cell self-renewal are known to involve the PI3K-AKT-mTOR signaling pathway. The pathway has been shown to be important regulator of NSC functions including self-renewal, neurogenesis, and cell survival (Akbar et al., 2005). DHA has also been shown to activate this pathway in Neuro 2A neuroblastoma cells (Akbar 2005; Akbar and Kim 2002; Kim, et al. 2000). Therefore, we examined the effects of DHA on this pathway in SVZ-derived neurosphere cultures. We found that DHA significantly increased phospho-AKT activity (Figure 4D) and that inhibiting the pathway with the PI3K inhibitor, LY294002, completely ablates all of the positive effects of DHA on stem cell self-renewal (Figure 4E) and neurogenesis (Figure 4F-G) indicating that PI3K activation is required for these effects, although does not exclude contributions from other pathways.

### DHA supplementation enhances SVZ cell survival and the longevity of mouse and rat neurosphere cultures

Our results show that there is a significant increase in cell survival (Figure 5A) and reduced caspase 3/7 activation in DHA-supplemented adult mouse SVZ cultures relative to vehicle control media conditions (Figure 5B). However, inhibition of PI3K (LY294002) ablates both the increase in cell viability (Figure 5A) and the decrease in caspase activation (Figure 5B), indicating that the PI3K pathway is required for the pro-survival effects of DHA treatment. It is well established that human and mouse SVZ neurosphere cultures derived from various regions of the embryonic CNS will expand in neurosphere culture for many months (Weiss et al., 1996; McKay, 1997; Palmer et al., 1997; Svendsen et al., 1998). In contrast, those derived from rat have a very limited capacity for exponential growth and undergo stem cell and culture senescence after only 4 or 5 passages (Svendsen et al., 1997). Given the positive effects of DHA on stem cell self-renewal and the subsequent rejuvenation of SVZ proliferation in vivo that we had observed, we tested if DHA could overcome this premature senescence in rat cultures. Our results demonstrate that DHA supplementation significantly enhanced the expansion of the rat cultures over control, DHA-depleted conditions from passage 2 onwards (Figure 5C). After passage two, there was a significant treatment effect increasing the number of cells at every passage. In addition, the fatty acid treatment increased the longevity of the rat cortical cultures from 4 passages in controls to 8 passages and beyond for the treated cells. Consequently, the cumulative number of cells generated over 8 passages was increased by approximately 280-400% over controls. However, despite this enhancement in cell expansion, the supplemented cultures still display a progressive decline in growth over time. Therefore, fatty acid supplementation appears to delay but does not fully overcome the onset of senescence in rat neurosphere cultures, suggesting that some studies which have examined DHA effects in rats and rat cultures may not be entirely reflective of the effects of DHA in mouse and human cells.

**Figure 5.**
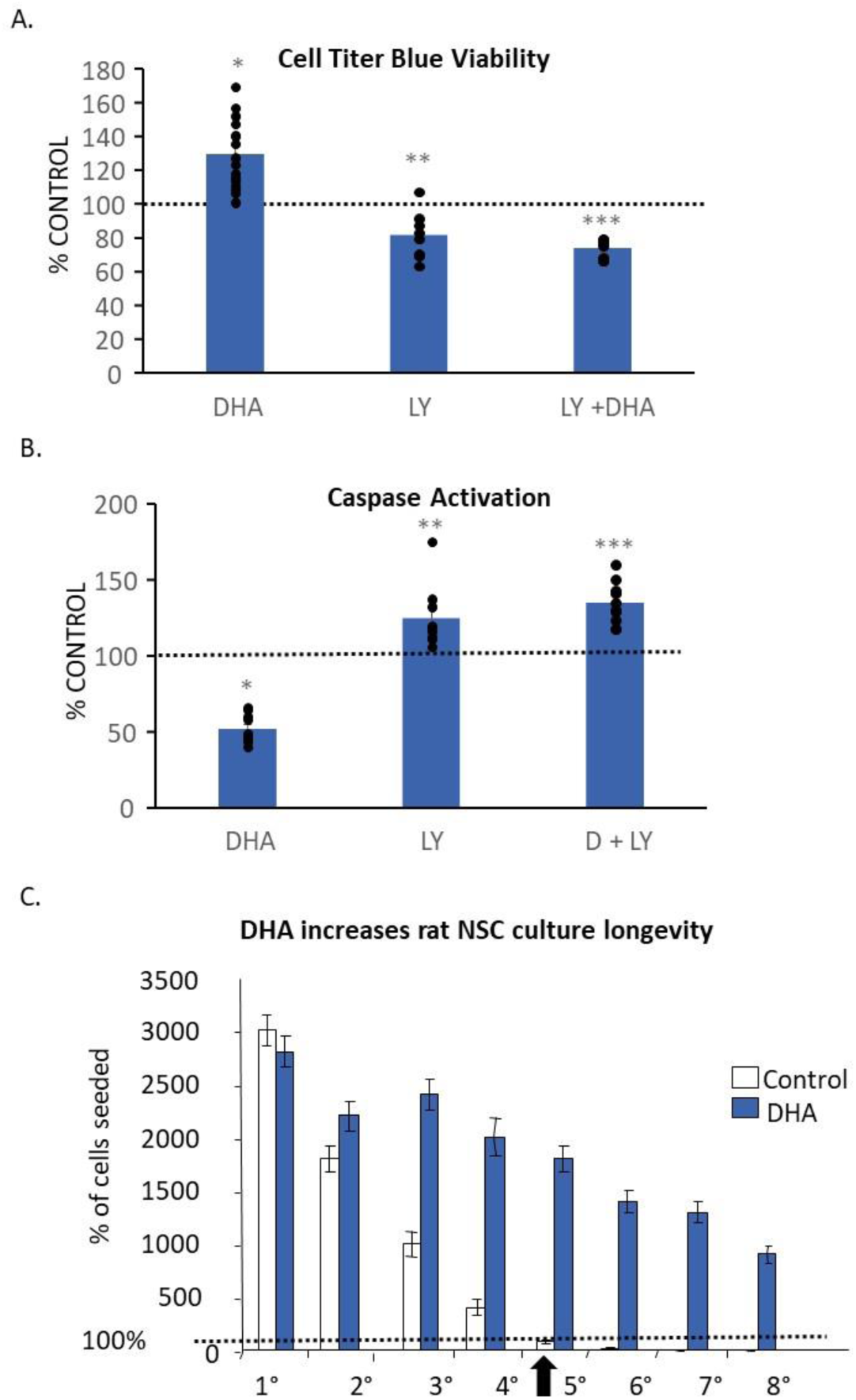
The positive effects of DHA on adult SVZ cell viability is mediated through the PI3K signaling pathway and increases the longevity of senescent adult rat SVZ cultures. (**A**) DHA supplementation of adult mouse SVZ cell cultures over 4 passages produces a significant increase in the number of viable cells in the culture compared to control conditions (P<0.001 Anova, post hoc Bonferroni *p=4.65×10^−6^, **p=0.028, ***p=4.03×10^−8^; n=16/grp) but inhibition of the PI3K pathway with the LY294002 inhibitor ablates that positive effect and diminishes culture viability (**p<0.05; n=8/grp); (**B**) the positive effects of DHA supplementation of adult mouse SVZ cultures can also be seen by its effects of caspase activity. DHA significantly decreased chemiluminescent-detected caspase activity in a Promega Caspase-glo plate assay and this effect was also ablated by PI3K inhibition with LY294002 (P<0.001; Anova post hoc Bonferroni *p=2.49×10^−5^, **p=0.019, ***p= 1.02×10^−3^); n=8/grp); (**C**) Adult rat neurosphere cultures are known to have a limited longevity in culture before they senesce and can no longer be passaged. We found that this occurs around serial passage 5 in our control conditions but the number of passages and therefore the total number of neural stem and progenitor cells produced by the cultures at each passage was extended out to at least passage 8 with DHA supplementation. Data shown are mean +/- SEM.

### DHA supplementation may increase the number of SVZ neural stem cells via an EGF-stimulated trans-differentiation of transit amplifying progenitors into multipotent stem cells

Given our results showing increased EGF sensitivity and our surprising findings that DHA was specifically enriching the stem cell population in the SVZ without causing changes to the overall proliferation rate of the cultures led us to examine the potential role of DHA in the phenomenon of neural stem cell trans-differentiation. It has been shown by us and by others that acute EGF exposure can increase SVZ stem cell numbers via a reversion of SVZ transit-amplifying cells back to a more immature stem cell phenotype (Thompson, et al. 2014; Doetsch et al., 2002). To test if this might contribute to the DHA-stimulated increases in the neural stem cell population that we observed with DHA treatments, we took freshly isolated SVZ cells and split them into treatment groups for a 4-hour incubation with and without EGF and with and without DHA in the culture media (see figure 6A). The time frame for this treatment is considerably less than the typical 12 to 24-hour cell cycle time of SVZ stem and progenitor cells so that changes in the relative numbers of stem cells in the cultures are not due to the proliferative production of new stem cell progeny. We found that within this acute time frame the freshly isolated SVZ cells that were split into different treatment groups had a small but significant 20% enrichment in the number of EGFR+ GFAP+ stem cells only when cultured with both EGF and DHA together (figure 6B). This demonstrates the potential for the increased EGFR sensitivity caused by DHA uptake combined with EGF stimulation to promote the production of new EGFR+GFAP+ stem cells from the EGFR+ transit amplifying progenitor population via a trans-differentiation back into a stem cell state. This could be sufficient to restore the SVZ stem cell population and improve neurogenesis without having a direct stimulatory effect on the rate of proliferation in the SVZ niche.

**Figure 6.**
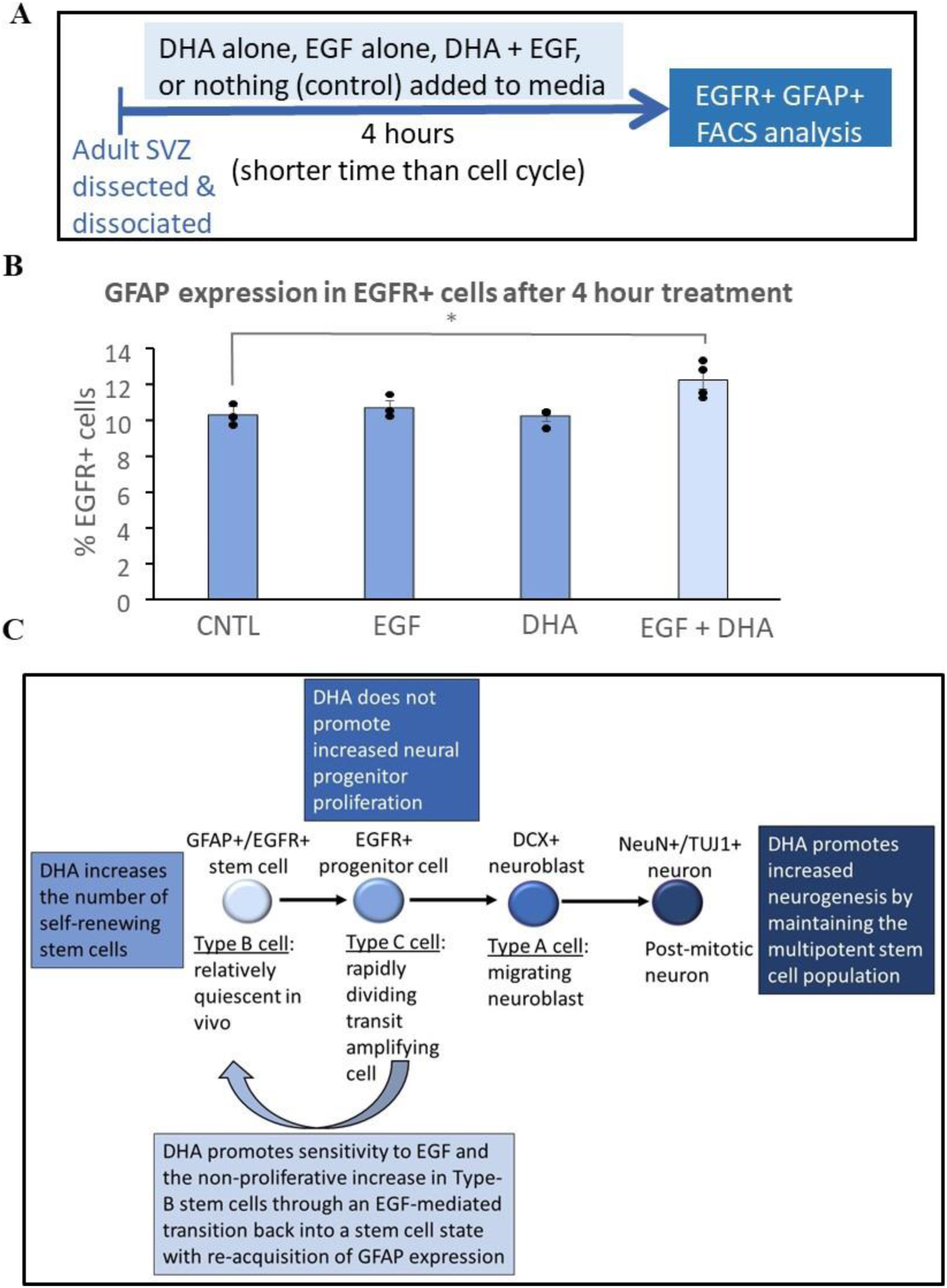
DHA may increase adult SVZ neural stem numbers and promote neurogenesis through an EGF-mediated trans-differentiation rather than proliferative process. (**A**) Representative timeline of the experimental paradigm showing how the effects of DHA on neural stem cell number following an highly acute treatment were examined. Adult mouse SVZ cells acutely dissociated from the brain were treated with DHA alone, EGF alone, or DHA and EGF combined for only 4 hours which is considerably shorter than the cell cycle time for these cells. Following this treatment FACS acquisition was used to assess the number of double-positive EGFR+ and GFAP+ cells (a known marker combination for significant neural stem cell enrichment); (**B**) FACS acquisition demonstrate that there is a small but significant (P=0.003 Anova post hoc Bonferroni *p=7×10^−3^; n=4 samples; 4 mice pooled per sample) increase in the number of EGFR+GFAP+ stem cells in the DHA and EGF treated group within a time frame that cell proliferation would not be able to significantly contribute to changes in cell numbers; (**C**) a summary diagram of our hypothesis based on these experiments of a potential mechanism for DHA to rescue SVZ function and neurogenesis in the aging brain. It illustrates effects of a non-proliferative increase in neural stem cell number in response to increased sensitivity to EGF stimulation that we have demonstrated which would then lead to an increase in the number of all proliferating progenitors in the SVZ which contribute to olfactory bulb neurogenesis (though not to any increases in the *rate* of proliferation). Thus, in this paradigm even a very small increase in the number of neural stem cells can produce a larger increase/rejuvenation in the number of their proliferative progeny (the EGFR+ transit amplifying progenitors) which are neurogenic. This unusual, non-proliferative mechanism could account for the positive cellular and behavior effects of DHA that we have observed.

## Discussion

During ageing, the rate of neurogenesis in both the SGZ and SVZ declines significantly (Kuhn et al., 1996; Enwere, et al., 2004; Hamilton et al., 2013). DHA enrichment in the brain has been shown to significantly increase neurogenesis in the young adult hippocampus (Dyall, et al., 2010; He et al., 2009; Janssen, et al., 2015) and to reverse age-related decline in hippocampal neurogenesis, neurite outgrowth, and memory function in aged mice (Cutuli, et al. 2014; Tokuda, et al., 2014; Hashimoto, et al., 2015; Calderon, et al., 2004; He, et al., 2009; Dyall, et al., 2010) but its effects on the aged SVZ neurogenic niche were largely unknown. We found that DHA has the ability to increase the number self-renewing neural stem cells and the neurogenic potential of the aged SVZ through a series of inter-related effects, possibly initiated by increased sensitivity to EGF receptor activation and elevated PI3K/AKT activity. An increase in the generation and survival of functional neural stem cells in response to DHA treatment also leads to a rejuvenation in the population of proliferative progenitors (though not in the rate of their proliferation) and neurogenesis in the aged SVZ. Recent studies on age-related changes to the SVZ neural stem cell niche have also suggested that the neural stem cells themselves have the capacity to be rejuvenated and reproduce younger levels of neurogenic function if the microenvironment of the niche itself is changed (i.e., reducing inflammation) (Kalamakis, et al., 2019). We have similarly shown that aged SVZ stem cells retain the capacity to be stimulated by growth factors to partially restore normal levels of self-renewing stem cells and neurogenesis with DHA treatment. It is known that brain and blood DHA levels decrease with age and this decline may be associated with the onset of dementia (Bazan, et al., 2011; Mohajeri, et al., 2015; Schaefer, et al. 2006). Thus, our data suggest that an important mechanism that potentially underlies the positive effects on cognitive function that previous clinical studies on dementia have observed with DHA treatment (Cole and Frautschy, 2010; Salem, et al., 2015; Morris, et al. 2003) may be its ability to restore the responsiveness of cells to growth factor stimulation in stem and progenitor populations which leads to improved adult neurogenesis. Unlike in the hippocampus, where DHA is known to have positive effects on learning and memory (Yurko-Mauro, et al., 2015; Sugasini, et al., 2017), the effects of DHA in the SVZ on olfactory function appear to result from selectively restoring the stem cell population rather than just generally enhancing proliferation in all dividing cells as has been reported in the dentate gyrus (He et al., 2009; Sakayori, et al., 2011). Thus, there may be significant differences in the mechanisms underlying the positive effects that DHA has on the aging brain between the different neurogenic niches.

The nervous system is enriched in the polyunsaturated fatty acids, arachidonic acid (AA, 20:4 n-6) and docosahexaenoic acid (DHA, 22:6 n-3), but neither can be synthesized de novo by mammals. Therefore, they and their precursors, linoleic and α-linolenic acid, are considered essential dietary fatty acids that must be transported into the brain. The conversion of α-linolenic acid to DHA is thought to occur primarily in the liver (Scott & Bazan 1989) and, to a lesser extent, by astrocytes and brain vascular endothelial cells which may secrete DHA into the cerebrospinal fluid for uptake by neurons. (Moore, et al., 1991, 2001; Balendiran, et. al. 2000). A major cellular uptake mechanism for DHA in the brain is mediated by brain fatty acid binding protein (B-FABP/FABP7/BLBP) which is a member of the FABP family of lipid chaperones with a very high binding affinity and specificity for DHA (Feng, et al., 1994; Xu, et al., 1996). The FABP7 isoform is only expressed in the central nervous system and is expressed at its highest levels during fetal development (Balendiran, et al., 2000; Storch and Corsico, 2008). FABP7 efficiently transports DHA and its precursor, α-linolenic acid, into all brain cells but it is most highly expressed in radial glia in the developing brain and in neural stem and progenitor cells in the adult brain (Balendiran, et al., 2000; Ebrahimi, et al., 2016; Giachino, et al. 2014; Matsumata, et al. 2012). In fact, the expression of the DHA-selective FABP7 protein is high enough in adult SVZ neural stem cells that it can be used in flow cytometry to selectively enrich for neural stem cells (Yun, et al., 2012). This close and selective association suggests a potentially important but unknown role for DHA in stem cell function. Our data supports this idea of stem cell-specific effects in the SVZ neurogenic niche by showing that the number of self-renewing stem cells, but not overall proliferation rates in all progenitors, is enhanced by DHA and that stem cell-enriched cultures using different prospective stem cell markers, including a FABP7-specific dye, are responsive to DHA treatment while stem cell-depleted cultures were non-responsive.

The effects of essential fatty acid deficits and supplementation have been widely studied in human health. It has been generally found to be the case that the n-6 and n-3 essential fatty acids have somewhat opposite biological effects with n-6 FAs being pro-inflammatory whilst n-3 FAs are anti-inflammatory (Galli and Calder, 2009; Mukherjee et al., 2004; Layé, et al., 2018) though they have many other functional effects beyond these. However, despite the clear evidence for the beneficial actions of fatty acids and their requirement by neurons, the underlying mechanisms for their effects on other cellular populations like stem and progenitor cells have been less well understood. The effects of DHA on different brain cell populations has been most extensively studied in the rat hippocampus. Several studies have investigated the effects of DHA on hippocampal progenitor populations and hippocampal function and found that it promotes neuronal differentiation and survival through a number of mechanisms including increasing retinoid-X receptor and pro-neuronal bHLH gene expression (Calderon and Kim, 2007; Katakura, et al., 2009; Kim et al. 2000; Dyall et al., 2010), it promotes progenitor proliferation in disease models (da Costa, et al., 2010; Borsini et al., 2017; Fiol-deRoque, et al., 2013), and it has a functional impact on learning and memory (Lozada et al., 2017; He et al., 2009). Our observations of SVZ cells have found the effects of DHA treatment on progenitor populations may differ significantly from the SGZ, such as a lack of a direct effect on the proliferation rate. However, despite potentially different mechanisms of action the functional effect of DHA that both niches share is enhanced neurogenesis and brain functions (learning, memory, olfaction, etc).

Several cellular studies have demonstrated that n-3 FAs, like DHA, can enhance neuronal differentiation, neurite outgrowth, and survival in cells from N2A neuroblastomas, mouse ES cultures, rat embryonic cortical cultures, and in the adult hippocampus of rats and FAT-1 transgenic mice (Kawakita et al., 2006; Cutuli, et al. 2014; He et al., 2009; Cao et al., 2005; Calderon & Kim, 2004; Akbar et al., 2005; Marszalek & Lodish, 2005; Katakura et al., 2009) but n-6 FAs, like arachidonic acid, do not (Katakura, et al., 2013). It has been observed that DHA treatment has anti-apoptotic effects following cell membrane incorporation and stimulation of phosphatidylserine synthesis on neural-like N2A and PC12 cells in culture (Kim et al., 2000, 2010) and in vivo in rat hippocampus and cortex providing neuroprotection from stressors such as ethanol exposure, inflammation-induced oxidative stress, hypoxia-ischemia, and traumatic injury (Ajami, et al., 2011; Zhang, et al., 2014; Paterniti et al., 2014; Williams et al., 2013; Berman, et al., 2009; Tajuddin, et al., 2014). Several studies have shown that some of these effects are due to the ability of DHA to be a positive modulator of the PI3K/AKT/mTOR signaling pathway (Akbar, et al., 2005; Akbar and Kim 2002; Kim, et al. 2000). We have found a similar stimulation of the PI3K/AKT/mTOR pathway in adult SVZ cells by DHA treatment which appears to be the primary signaling pathway by which DHA contributes to increased survival in the mouse cultures and increased longevity achieved in rat cultures which normally stop growing after only a few passages in vitro (Svendsen et al., 1997). Given the known association of the PI3K signaling pathway with neural stem cell self-renewal (Groszer et al., 2001, 2006), it seems likely to contribute to the increased number of multipotent neural stem cells that we observe with DHA supplementation. For example, in mutant mice with a conditional genetic deletion of the tumor suppressor PTEN, a negative regulator of PI3K, there is a large increase in pathway activation and subsequent increases in SVZ neural stem cell self-renewal and neurogenesis resulting in olfactory bulb overgrowth (Gregorian et al., 2009). In our current studies we observed that PI3K inhibition in SVZ cultures blocks all of the positive effects of DHA on clonal neurosphere formation, survival, and neurogenesis suggesting that DHA does primarily exert its effects on multiple cell functions via this signaling pathway. However, it was not initially clear how this pathway activation may be selectively affecting SVZ stem cell self-renewal without also enhancing overall neural progenitor proliferation rates. Our data on the effects of DHA on EGF receptor activity and expression led us to develop and test a hypothesis of a non-proliferative trans-differentiation mechanism that can result from increased EGFR sensitivity which could explain these unique effects of DHA on the SVZ.

DHA has been shown to be primarily incorporated into membrane phospholipids altering membrane fluidity and the expression and function of many membrane-associated proteins which can affect things like receptor mediated signal transduction and ion channel activity (Rapoport 2001; Leaf et al., 2002; Stillwell and Wassall, 2003; Stillwell, et al. 2005; Turk et al., 2012; Bazan, 2005).For example, DHA can also bind nuclear receptors (i.e., Peroxisome Proliferator-Activated Receptors-PPARs and Retinoid Acid X Receptors-RXRs) which regulate transcription (Lengqvist, et al., 2004; de Urquiza, et al., 2000; Kothapalli et al., 2007). These mechanisms, working separately or in concert, contribute to DHA’s functional effects on the brain. Importantly, it has been found that DHA-supplemented neural stem and progenitor cultures derived from embryonic rat cortex have an altered membrane lipid profile which specifically affects the expression of EGF receptor and EGFR phosphorylation (activation) following EGF exposure (Langelier et al., 2010). In line with these findings, our data show that DHA treatment of SVZ stem and progenitor cultures produces an enhanced functional sensitivity (stem cell self-renewal) and EGFR phosphorylation in response to EGF stimulation in vitro. Though it is well established that EGFR stimulation activates the PI3K/AKT signaling pathway (Wee, et al., 2017), it has now been shown by the Doetsch lab that this activation can also contribute to a reversion of SVZ transit-amplifying cells back to a more immature stem cell phenotype rather than promoting a proliferative response (Doetsch et al., 2002). We have also observed this phenomenon to occur with EGFR+ SVZ transit amplifying progenitors following traumatic brain injury (Thompson, et al. 2014) which is known to produce elevated brain DHA levels (Guo, et al., 2017). Our current data suggest that the DHA-enhanced sensitivity to EGF stimulation that we have observed in the adult SVZ cells could potentially contribute to a similar reversal of the normal lineage progression of GFAP+/EGFR+ stem cells into EGFR+ transit-amplifying cells. Our results from this experiment suggest that EGFR+ transit amplifying progenitors respond to combined DHA and EGF stimulation to produce a non-proliferative, de-differentiation expansion of the SVZ stem cell population (see summary figure 6C). If this is the primary functional effect of DHA on the SVZ stem cell population, then it does not appear to have any pro-proliferative effects on either the progenitor or stem cell populations in the adult SVZ. Instead, the restoration of normal (younger) adult levels of proliferation and neurogenesis and olfactory function that we have observed in aged mice fed a diet enriched with DHA may result from this restoration in stem cell numbers within the SVZ which would then lead to normal (younger), but not enhanced, rates of progenitor proliferation and neurogenesis.

Finally, these observations contribute to the current consensus that the stem cells within the aging SVZ niche retain their intrinsic ability to self-renew and produce proliferative, neurogenic progeny which can be restored through the manipulation of the niche environment, in this case with DHA enrichment. Thus, despite the recent debate over whether any post-natal SVZ neurogenesis occurs in humans, our data suggest that this deficit could potentially be overcome through DHA enrichment or enhancing EGF receptor activation and PI3K signaling activity in the human SVZ niche, which has hopeful implications for countering age-related cognitive decline and for promoting endogenous brain repair after injury.

## Acknowledgements

The authors would like to gratefully acknowledge Martek Biosciences for providing the DHASCO oil to produce the custom rodent feed for our experiments. We would also like to thank Professor Chang Young-Tae for generously providing his CDr3 FABP7 dye before it became commercially available.

